# Nutrient-dependent allometric plasticity in a male-diphenic mite

**DOI:** 10.1101/2021.06.14.448383

**Authors:** Flor T. Rhebergen, Kathryn A. Stewart, Isabel M. Smallegange

## Abstract

Male secondary sexual traits often scale allometrically with body size. These allometries can be variable within species, and may shift depending on environmental conditions such as food quality. Such allometric plasticity has been hypothesized to initiate local adaptation and evolutionary diversification of scaling relationships, but is under-recorded, and its eco-evolutionary effects are not well understood. Here, we test for allometric plasticity in the bulb mite (*Rhizoglyphus robini*) in which large males tend to develop as armed adult fighters with thickened third legs, while small males become adult scramblers without thickened legs. We first examined the ontogenetic timing for size- and growth-dependent male morph determination, using experimentally amplified fluctuations in growth rate throughout juvenile development. Having established that somatic growth and body size determine male morph expression immediately before metamorphosis, we examined whether the relationship between adult male morph and size at metamorphosis shifts with food quality. We found that the threshold body size for male morph expression shifts towards lower values with deteriorating food quality, confirming food-dependent allometric plasticity. Such allometric plasticity may allow populations to track prevailing nutritional conditions, potentially facilitating rapid evolution of allometric scaling relationships.

## Introduction

Male secondary sexual traits are often conspicuously sensitive to variation in juvenile food quality (Emlen & Nijhout, 2000; Kodric-Brown *et al*., 2006; Lavine *et al*., 2015). This nutrition-sensitive plasticity generally manifests itself in hyper-allometric scaling of sexual traits with body size (Simmons & Tomkins, 1996; Kawano, 2000; McCullough *et al*., 2015) or even in discrete alternative reproductive phenotypes (Emlen *et al*., 2005; Smallegange, 2011). As a consequence, large individuals in a population can have enormous secondary sexual traits, while small individuals can be inconspicuously endowed. Scaling relationships can vary across time and space within species (Miller & Emlen, 2010). Such variation is sometimes due to genetic variation; different populations can differ in genetic background due to drift or local adaptation, resulting in variation in how body size relates to trait size (Buzatto *et al*., 2012). It is also possible that the scaling relationship itself exhibits plasticity in response to local environmental effects, a phenomenon that has been dubbed ‘allometric plasticity’ (Emlen, 1997; Moczek, 2002; Casasa & Moczek, 2019). Allometric plasticity can occur in response to food variation (Emlen, 1997; Moczek, 2002). In such cases, variation in nutrition not only causes variation in body size and therefore in sexual traits, but also variation in *how* organisms weigh the body size cue to develop sexual traits.

Food-dependent allometric plasticity has a significant role in driving ecological and evolutionary patterns in sexual traits. Ecologically, allometric plasticity causes divergence in scaling relationships among populations in different nutritional environments. Such inter-population variation is often interpreted as local adaptation (Tomkins & Brown, 2004) or speciation in progress (Kawano, 2002), but may instead reflect undetected allometric plasticity. Allometric plasticity can also obscure scaling relationships of interest within populations when environments are nutritionally heterogeneous on small spatial or temporal scales. Such intra-population variation in scaling can conceal condition-dependence, which is often inferred through static allometric scaling (Bonduriansky, 2007; Eberhard *et al*., 2018), and leave the impression that the association of sexual trait variation with body size is much weaker than it actually is. Evolutionarily, allometric plasticity is hypothesized to be an important first step in adaptive divergence of scaling relationships in novel (nutritional) environments (Casasa & Moczek, 2019). Genetic accommodation of ancestrally plastic scaling relationships can allow rapid adaptive evolution, if allometric plasticity in the novel environment coincides with release of selectable cryptic genetic variation, and/or if the novel environment persists for long enough to allow genetic mutations to accumulate and canalize the induced scaling relationship (West-Eberhard, 2003; Pfennig *et al*., 2010; Moczek *et al*., 2011; Casasa & Moczek, 2019).

Despite the hypothesized ecological and evolutionary importance of food-dependent allometric plasticity, it has been demonstrated only in a few species with continuous positive sexual trait allometries (Emlen, 1997; Knell *et al*., 1999; Moczek, 2002). In the majority of species with allometrically scaling sexual traits, such allometries have been measured in (non-)randomly sampled individuals from the natural environment, without information on environmental circumstances during development beyond body size (e.g. Kawano, 2000; Buzatto *et al*., 2014; McCullough *et al*., 2015). Therefore, to assess whether food-dependent allometric plasticity affects the eco-evolutionary dynamics of sexually selected condition-dependent traits, it is first and foremost necessary to assess whether such allometric plasticity occurs in other well-known model systems for the evolution of condition-dependent sexually selected traits.

Here, we tested whether food-dependent allometric plasticity is present in a species with discrete, condition-dependent, alternative male morphs: the bulb mite *Rhizoglyphus robini* (Radwan, 1995; Smallegange, 2011). In *R. robini*, large males develop the adult ‘fighter’ morph, which has a proportionally thickened third leg pair with dagger-like claws that it uses in fights to kill rival males (Radwan *et al*., 2000; Fig. 1a). Small males tend to develop the adult ‘scrambler’ morph, which has a normal, female-like, third leg pair and a modest motility advantage (Tomkins *et al*., 2011; Fig. 1b). Although it is well established that the alternative morphs in *R. robini* are associated with variation in body size, and thus represent a discontinuous allometric scaling relationship (Smallegange, 2011; Leigh & Smallegange, 2014; Smallegange & Deere, 2014), the correlation is noisy. A large range of overlap in body size exists between juveniles developing as fighters and juveniles developing as scramblers (Smallegange, 2011; Stewart *et al*., 2018). We hypothesize that these observations are, at least partly, explained by food-dependent plasticity in how body size scales with male morph expression. Such allometric plasticity, if present, could facilitate the repeatedly observed rapid evolution of how male morph expression relates to body size in *Rhizoglyphus* mites (Tomkins *et al*., 2011; Smallegange & Deere, 2014; Casasa & Moczek, 2019).

**Figure 1.**
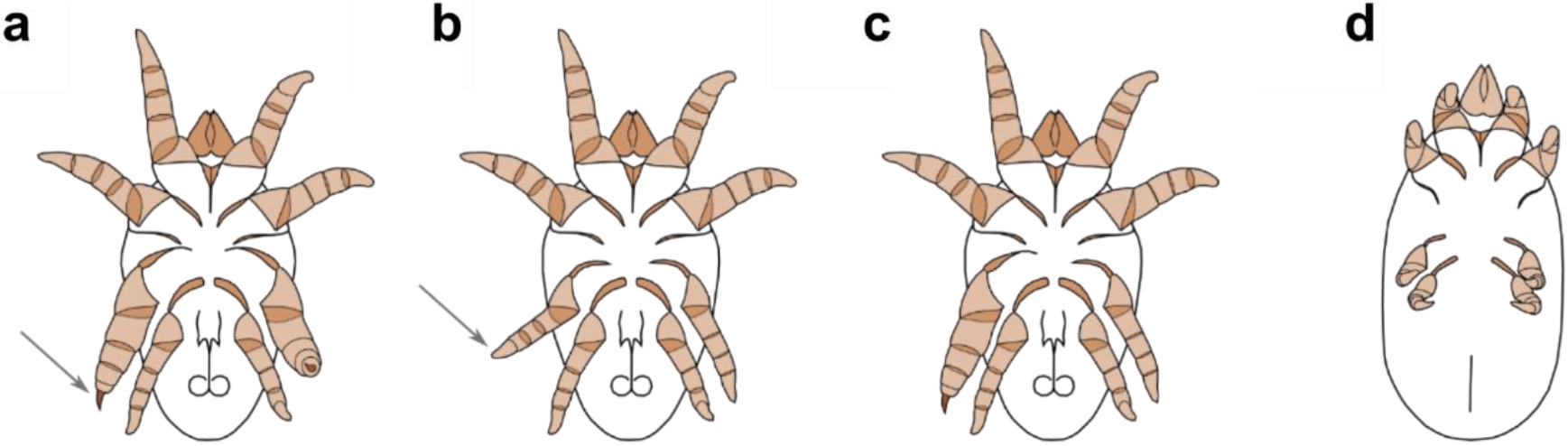
*Rhizoglyphus robini* morphology, ventral view. **a**, Adult fighter male. **b**, Adult scrambler male. **c**, Adult intermorph male. **d**, Quiescent tritonymph stage, during which metamorphosis and fighter leg development takes place. Fighters and scramblers differ in the width of the third leg and in the shape of the third tarsal claw (grey arrows).

To test the hypothesis that condition-dependent male morph expression is associated with food-dependent allometric plasticity, we first identified the sensitive ontogenetic stage at which male body size (or a physiological correlate of it) cues male morph expression. Assuming that growth rate and/or body size explain morph expression best during this sensitive stage, we exposed males to different fluctuating patterns of food availability during ontogeny, creating random variation in growth trajectories from larva to adulthood. We then assessed whether the resulting variation in male morph expression was fully explained by variation in growth rate and/or body size, at different time frames in ontogeny. Having identified the sensitive stage of morph determination, we then tested whether the allometry of adult third leg width and body size at the sensitive stage of morph determination shifts with food quality. We find that adult male morph expression is cued by growth rate and body size immediately before metamorphosis, and that this threshold body size at metamorphosis shifts towards smaller values with deteriorating food quality. Such allometric plasticity in *R. robini* can allow the body size threshold for male morph expression to adaptively track nutritional conditions and resulting shifts in the body size distribution.

## Materials and Methods

### Study species and general procedures

*Rhizoglyphus robini* is a cosmopolitan agricultural pest mite on numerous crops and forms dense populations on subterraneous plant structures (Díaz *et al*., 2000). The populations in this study were derived from *R. robini* samples that were collected from flower bulbs in storage rooms in Noord-Holland, The Netherlands, in December 2010. In favourable environments, *R. robini* has four morphological life history stages (larva, protonymph, tritonymph, then adult), interspersed with quiescent phases during which individuals do not move or eat, but undergo incomplete metamorphosis to enter the next stage (Fig. 1d).

We keep our *R. robini* populations in an unlit climate cabinet (25 °C; >90% humidity), in plastic containers (10 × 10 × 2.5 cm) with a plaster of Paris substrate. Twice a week, we clean one sixth of the substrate by removing all organic matter and add fresh water and food. We keep half of our populations (four containers) on a diet of baker’s yeast and the other half on a diet of oats. Egg-to-egg time averages 11 days on yeast and 13 days on oats when food is *ad libitum*. Otherwise, mites from both diets are similar; no morphological or behavioural differences have been recorded between our yeast-fed and oats-fed populations (unpublished data). Populations on the same diet are mixed every six months to maintain genetic homogeneity.

In the experiments, individual mites were kept in individual plastic tubes (height 5 cm, diameter 1.5 cm) with a moistened plaster substrate. The tubes were sealed with fine mesh, kept in place by a plastic lid and placed in an unlit climate cabinet (25 °C; >90% humidity) throughout the experiment. We avoided bias due to genetic variation by randomly sampling the parental generation from our stocks, and allocating one offspring per parental pair to each experimental treatment – an isoline approach to test several genotypes explicitly would have been logistically infeasible. Mites were photographed ventrally and measured to the nearest 0.001 µm using a Zeiss Axiocam 105 color camera mounted on a Zeiss Stemi 200-C stereomicroscope, and ZEN 2 (Blue edition) software.

### Experiment 1: When during ontogeny is male morph expression cued by body size and/or growth?

The mites in this experiment were reared on a diet of oats rather than yeast because development is slightly slower on oats than on yeast, increasing our power to find the critical developmental period in which morph expression is determined. The parental generation was formed by collecting 324 individual *R. robini* larvae from the oats-fed stock populations. The parental larvae were reared to adulthood on a diet of *ad libitum* oats in individual tubes. Adult females and males (90.1% fighters, 8.3% scramblers, 1.6% intermorphs, i.e. males that were fighter on one side and scrambler on the other (Fig. 1c)) were randomly paired to produce the focal generation. Pairs were kept in isolation in separate tubes for three days and were fed *ad libitum* oats. From each parental pair a single egg was collected and individually isolated. These individuals were reared to adulthood on a variable (n = 81) or a constant (n = 21) diet of powdered oats to obtain a wide range of growth trajectories. Focal individuals that were reared on a variable diet were transferred daily to a fresh tube that either did or did not contain food. The probability of changing to a different environment (i.e., moving from a food to a non-food tube, and vice versa) was set to 0.8 (n = 27), 0.5 (n = 27) or 0.2 (n = 27). Each individual was photographed and measured at 24-hour intervals to record growth trajectory. We measured idiosoma width and recorded morph expression upon maturation.

To test whether male morph expression was a response to food uptake and growth at the final moments of the tritonymph stage, versus earlier during juvenile growth, we constructed multiple logistic regression models. For each of six developmental stages, we constructed a generalized linear model (GLM) with a logit link function, with morph expression (fighter or scrambler) as response variable, and growth during that stage and size when entering that stage as explanatory variables. The six developmental stages we distinguished were the first and second half of the larval, protonymph and tritonymph stages (Table 1). We defined ‘size when entering the first half of the larva stage’ as the length of the egg, ‘size when entering the second half of the larva, protonymph or tritonymph stage’ as the idiosoma width of an individual when it had spent half of its total time in its respective stage, and ‘size when entering the first half of the protonymph or tritonymph stage’ as the idiosoma width of the quiescent larva or protonymph, respectively, ‘growth during a particular stage’ as the difference in idiosoma width when entering a particular stage (early/late larva, early/late protonymph or early/late tritonymph) and when entering the following stage. ‘Growth during the early larval stage’ was not available because we did not have idiosoma measurements at hatching. Therefore, we included idiosoma width at the end of the first half of the larval stage, rather than growth during the first half of the larval stage, as the first explanatory variable for the early-larva logistic regression model. We assessed the performance of the logistic regression models for the different stages by comparing AICc values (Akaike Information Criterion, corrected for small sample size) and calculating the percentage deviance explained. We tested whether the models performed better than chance by comparing the fitted models to null models using LRT. All analyses were done in R v. 3.4.4 (R Core Team, 2018), including packages *stats, dplyr* (Wickham *et al*., 2018) and *ggplot2* (Wickham *et al*., 2019).

**Table 1.** Logistic regression models with male morph expression (response variable), and size at the beginning of a juvenile stage and growth during that stage (explanatory variables), for each juvenile stage. The difference in deviance between the fitted models and null models without any explanatory variables is χ^2^-distributed under the null hypothesis, with 2 degrees of freedom.

| Juvenile stage | AIC<sub>c</sub> | Null deviance | Residual deviance | Difference in deviance ($\chi^2$, df = 2) | P-value | % deviance explained |
| --- | --- | --- | --- | --- | --- | --- |
| Early larva | 36.39 | 30.885 (31 df) | 29.533 | 1.352 | 0.509 | 4.4 % |
| Late larva | 34.30 | 30.885 (31 df) | 27.439 | 3.446 | 0.179 | 11.2 % |
| Early protonymph | 30.82 | 29.569 (28 df<sup>†</sup>) | 23.860 | 5.709 | 0.058 | 19.3 % |
| Late protonymph | 32.97 | 29.569 (28 df<sup>†</sup>) | 26.012 | 3.557 | 0.169 | 12.0 % |
| Early tritonymph | 32.31 | 30.885 (31 df) | 25.455 | 5.43 | 0.066 | 17.6 % |
| *Late tritonymph* | *16.37* | *30.885 (31 df)* | *9.516* | *21.369* | *<0.001* | *69.2 %* |

† Three fighter males passed the protonymph stage very quickly precluding multiple protonymph measurements. These males were removed from protonymph models.

### Experiment 2: Is size-dependent male morph expression subject to allometric plasticity?

The parental generation was formed by individually isolating 200 larvae from the yeast-fed stock populations, in four experimental blocks of 50 larvae each. These larvae were reared to adulthood on a diet of *ad libitum* yeast in individual tubes. Per block, 15-24 adult males and females were randomly paired in a plastic tube with *ad libitum* yeast to produce the focal generation. Three days after pairing, eggs of each parental pair were transferred to a clean individual tube without food. Within 12 hours after hatching, larvae were transferred to experimental tubes. Each parental pair contributed a single larva to each food treatment.

The focal individuals were individually isolated and reared to adulthood on quartered ∼23 mm^2^ circular disks of filter paper that were soaked for 20 seconds in 10 μL solution of yeast in water. Individuals were subjected to five food treatments, which represented a 2.5-fold dilution series (yeast solved in water): 40.0 mg/mL (n = 69), 16.0 mg/mL (n = 40), 6.40 mg/mL (n = 40), 2.56 mg/mL (n = 69) and 1.02 mg/mL (n = 40). Experimental blocks 1 and 4 contained all treatments, but due to logistical constraints, blocks 2 and 3 only contained the 40.0 mg/mL and 2.56 mg/mL treatments. Every two days, the substrate of each tube was cleaned with a moist brush and the yeast-soaked disk was replaced with a fresh disk. After individuals had entered the quiescent tritonymph stage, the disk was removed; adults emerged in an empty tube.

Each individual was photographed and measured in the quiescent tritonymph (Fig. 1d) and adult stage (Fig. 1a-c). We measured the width of the idiosoma (the white bulbous posterior part of the body) in the quiescent tritonymphs and adults (st. dev. = 3.3 µm; Supplementary Information) and the width of both third leg femurs in the adults (st. dev. = 1.1 µm), and recorded male morph expression.

To investigate the effect of food quality and QTIW on the probability of fighter expression, we constructed a logistic regression model with fighter expression as response variable, QTIW, food quality, the interaction between QTIW and food quality and block as fixed explanatory variables and parent ID as random intercept. Four intermorph males were excluded from this analysis. We centered QTIW around the overall mean QTIW value, so that the intercept represented the log-odds of fighter expression at the average QTIW. We tested significance of explanatory variables using LRT.

## Results

### Experiment 1: When during ontogeny is male morph expression cued by body size and/or growth?

We were able to follow the developmental trajectories of 69 focal individuals from egg to adulthood (67.6% of total isolated eggs). The individuals that we were not able to follow either died during experimentation or were accidentally lost during handling. Of these 69 individuals, 33 were males, 26 of which expressed the fighter morph. Adult morph expression was predicted by idiosoma growth and size in the last half of the tritonymph stage, immediately prior to metamorphosis (GLM: 69.2% deviance explained, χ^2^ = 21.37, df = 2, P < 0.001; Table 1), but not in the first half of the tritonymph stage, the first and second half of the protonymph stage, or the first and second half of the larval stage (Table 1). Specifically, males that were large at the beginning of the last half of the tritonymph stage had a higher probability of expressing the fighter morph than small males, as did males that grew relatively faster during the last half of the tritonymph stage. The scrambler morph was expressed only by males that were very small at the beginning of the last half of the tritonymph stage, or grew little during the last half of the tritonymph stage (Fig. 2a). Morph expression also did not depend on idiosoma growth and size in preceding juvenile stages prior to the second half of the tritonymph stage (Fig. 2b-f). For earlier stages, size-at-beginning-of-stage and growth-during-stage did not significantly explain variation in morph expression (Table 1).

**Figure 2.**
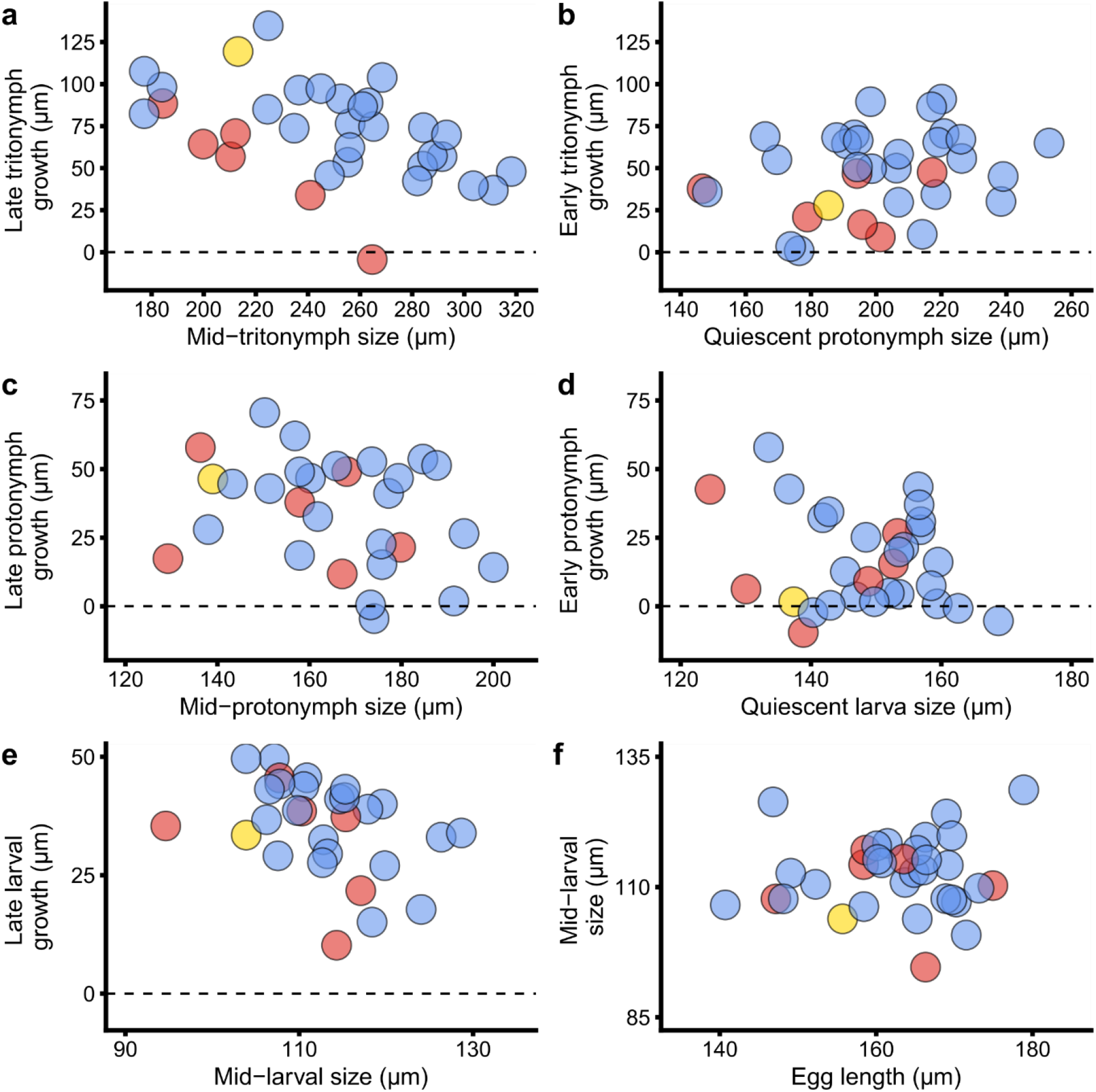
Male morph expression (blue: fighter, red: scrambler, yellow: intermorph) as a response to experimentally induced variation in size at the beginning of a particular juvenile developmental stage, and growth during that stage. **a**, Second half of the tritonymph stage. **b**, First half of the tritonymph stage. **c**, Second half of the protonymph stage. **d**, First half of the protonymph stage. **e**, Second half of the larval stage. **f**, First half of the larval stage.

### Experiment 2: Is size-dependent male morph expression subject to allometric plasticity?

We reared a total of 128 males to adulthood on filter disks soaked in a yeast concentration of 40.0 mg/mL (n = 35), 16.0 mg/mL (n = 21), 6.40 mg/mL (n = 12), 2.56 mg/mL (n = 37) and 1.02 mg/mL (n = 23). The interaction between QTIW and food quality was statistically significant (LRT: χ^2^ = 11.78, df = 4, P = 0.019), indicating that the effect of QTIW on the probability of fighter expression varied across food treatments. Upon visual inspection of the data, the significant interaction between QTIW and food quality was likely caused by two artefacts of limited sampling: (i) the effect of QTIW on fighter expression had vanished in 6.40 mg/mL. However, in this treatment we could sample only 12 males, four of which became scramblers and one expressing the intermorph phenotype; (ii) QTIW completely separated fighters (n = 27) from scramblers (n = 7) in 40.0 mg/mL. Therefore, we assumed the true interaction effect to be zero (constraining the slopes of the logistic regression curves to the same value) and proceeded to test the main effects of QTIW and food quality. The probability of fighter expression given the effect of food type increased significantly with QTIW (LRT: χ^2^ = 75.83, df = 1, P < 0.001; Fig. 3a-f). Food quality also had a highly significant effect on the probability of fighter expression given the effect of QTIW (LRT: χ^2^ = 44.49, df = 4, P < 0.001), showing a plastic body size threshold that scaled with food quality (Fig. 3a-f).

**Figure 3.**
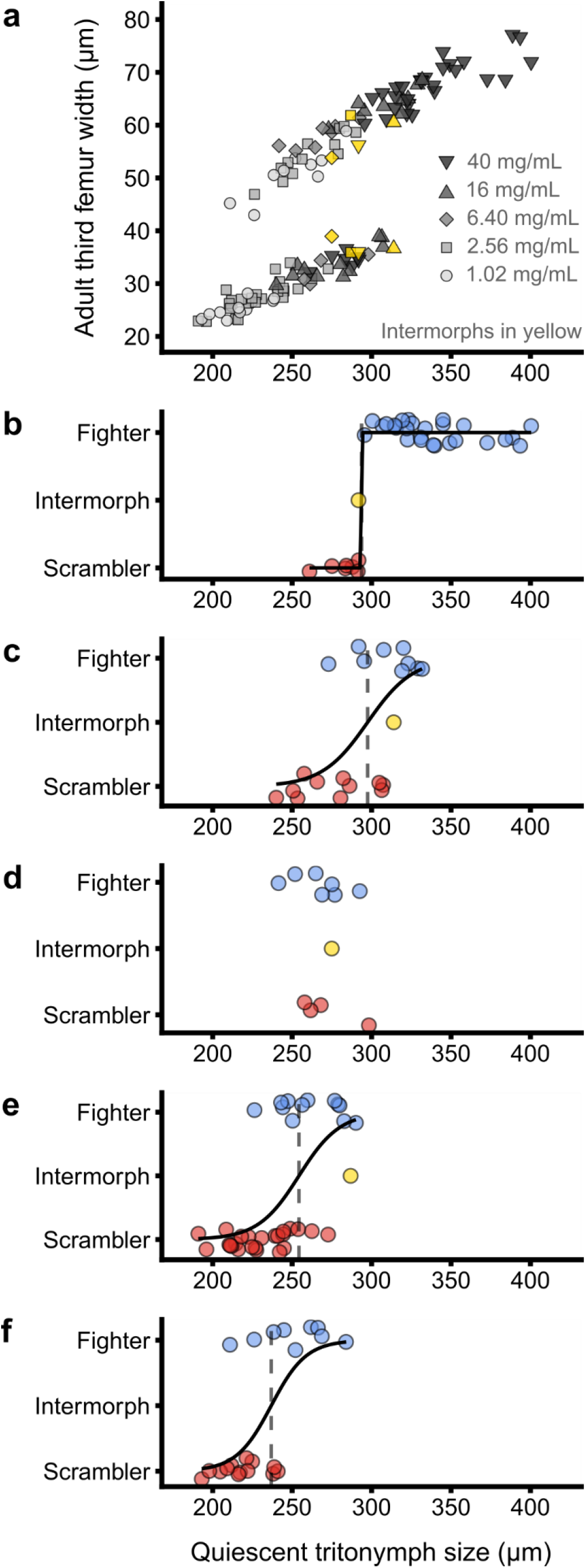
Condition-dependent enlarged fighter leg development shows food-dependent allometric plasticity. **a**, The allometric relationship of adult male third femur width (left and right leg averaged) and body size at metamorphosis. Colour and shape denote food quality (filter paper soaked in yeast solution; see legend). Intermorph males are shown in yellow; in this case, both third legs are shown. **b-f**, Male morph expression (blue: fighter, red: scrambler, yellow: intermorph) as a response to quiescent tritonymph body size, for males raised on a diet of 40.0 mg/mL yeast (**b**), 16.0 mg/mL yeast (**c**), 6.40 mg/mL yeast (**d**), 2.56 mg/mL yeast (**e**) and 1.02 mg/mL yeast (**f**). Solid black lines represent statistically significant (α = 0.05) logistic regression lines and denote the probability of fighter expression. Vertical dashed lines represent the quiescent tritonymph size threshold at which the probability of fighter expression is estimated to be 50%.

## Discussion

We hypothesized that food-dependent allometric plasticity underlies size-dependent male morph expression in *Rhizoglyphus robini*. If this is the case, the allometry of adult third leg width and body size at the developmental stage when male morph expression is cued should change depending on perceived food quality. We found that expression of the fighter phenotype is associated with body size and growth rate during the second half of the final juvenile stage before metamorphosis, but not earlier during ontogeny. This suggests that male morph expression is cued immediately before or during metamorphosis, in the quiescent tritonymph developmental stage. We then tested whether the allometry of quiescent tritonymph body size and the width of the adult third legs shifts with food quality. We found that the allometry did shift towards lower body sizes with a deterioration in food quality, confirming allometric plasticity in *R. robini* male weaponry.

The pattern of food-dependent allometric plasticity can emerge from different developmental mechanisms. Firstly, it is possible that male tritonymphs weigh and integrate multiple cues from the nutritional environment, to assess future mating chances and adaptively adjust development (Emlen, 1997). Hypothetically, nutrition and growth inform not only on a male’s own future physical strength, but also on the strength of potential rivals, provided that these rivals experience the same nutritional circumstances. Therefore, a male’s developmental decision to prepare for an aggressive mating tactic should depend not just on its own body size, but on its body size relative to the expected average, favouring adaptive allometric plasticity (Emlen, 1997).

A second explanation for the observed allometric plasticity is that males use only a single cue for fighter expression, which is not body size, but instead *correlates* with body size. If the association of the ‘true cue’ and body size depends on food quality, allometric plasticity will emerge. This could happen, for example, if male morph expression is cued by an individual’s resource budget during metamorphosis (Smallegange *et al*., 2019). Larger individuals will have larger resource budgets on average, but resource budgets are not expected to scale linearly with body size, because the energetic expenses associated with growth and metamorphosis also increase with size. Therefore, allometric plasticity could be a symptom of budget-dependent development of enlarged and expensive fighter legs.

Finally, apparent allometric plasticity could emerge if male morph expression and relative body size are both pleiotropically influenced by genes of large effect. If that is the case, allometries are effectively genetically determined, while nutrition simply shifts the genetically determined allometries with average body size. Male morph expression in *R. robini* is certainly heritable (Radwan, 1995, 2003; Skrzynecka & Radwan, 2016), and fighter expression has been shown to be associated with genetic diversity and potentially coadaptive gene complexes (Stewart *et al*., 2019) and the upregulation of genes involved in general metabolism (Joag *et al*., 2016). However, to our knowledge, there is currently no evidence suggesting that the link between male morph expression and growth rate is caused by pleiotropy, rather than developmental plasticity of leg development in response to growth, size, the resource budget, or food quality. Pleiotropic genetic control of growth and male morph expression seems improbable given the results of our first experiment. Nonetheless, it is likely that the proportion of growth variation explained by genetic effects is larger when the nutritional environment is stable over time (as in our experiment 2) than when the nutritional environment is strongly fluctuating (as in our experiment 1). Likewise, male morph expression may be largely genetically influenced in unfluctuating nutritional environments, and more strongly affected by nutrition in fluctuating nutritional environments. More work is needed to exclude the possibility that apparent allometric plasticity in fact reflects a genetically controlled allometry, driven by genes of large effect, pleiotropically affecting both growth and male morph expression. However, to our knowledge, such a genetically controlled allometry would be a previously unrecorded phenomenon in arthropods.

Allometric plasticity is hypothesized to be an important first step in the evolutionary diversification of allometric scaling relationships (Casasa & Moczek, 2019), and has been recorded in a few species with conspicuous sexual trait allometries. In the dung beetles *Onthophagus acuminatus* and *O. taurus*, the allometry of male horns and adult body size plastically shifts with food quality and average body size (Emlen, 1997; Moczek, 2002), so that the allometry tracks a shift in the body size distribution much like we describe for *R. robini*. These examples of allometric plasticity have been hypothesized to reflect adaptation to variation in the expected competitive environment, reflected by seasonally varying food (e.g. dung quality) (Emlen, 1997). Alternatively, they may reflect a developmental side-effect due to additional (but food-dependent) growth after secondary sexually selected traits have been cued by body size (Moczek, 2002). The opposite pattern of food-dependent allometric plasticity has also been recorded, where allometric plasticity does not track shifts in the body size distribution, but rather amplifies condition-dependence. For example, in the stalk-eyed fly *Diasemopsis aethiopica* and the armed beetle *Gnatocerus cornutus*, the allometric scaling of sexual trait size is lowered in individuals raised on poor-quality compared to individuals raised on good-quality food (Knell *et al*., 1999; Okada & Miyatake, 2010). Allometries can also change in response to seasonal change: in the stag beetle *Lucanus cervus*, the allometry of male mandibles and adult body size changes over the course of summer, so that males pupating later in the year develop smaller mandibles for their body size than males pupating early in the year (Hardersen *et al*., 2011). In this case, allometric plasticity is hypothesized to be adaptive due to time constraints on development in combination with allocation trade-offs: the mating season is short, so late-developing males should accelerate maturation at the expense of energetic investment in weaponry (Hardersen *et al*., 2011). Yet another form of allometric plasticity occurs in the mite species *Rhizoglyphus echinopus* and *Sancassania berlesei*, which exhibit the same size-dependent fighter-scrambler polyphenism as the closely related *R. robini*. In these species, but curiously not in *R. robini* (Radwan, 1995), fighter expression in large males is suppressed by pheromones from dense colonies, so that the allometry collapses into complete scrambler expression in dense colonies (Radwan, 1993, 2001). This response has been demonstrated to be adaptive in *S. berlesei*: in large groups, the fighter phenotype no longer yields higher mating success than the scrambler phenotype even for large males (Michalczyk *et al*., 2018), and instead becomes a liability (Radwan & Bogacz, 2000; Radwan *et al*., 2000).

The different forms of allometric plasticity showcase the developmental flexibility associated with allometric scaling, but also highlight that these apparently similar patterns may have evolved under different selection pressures. Therefore, to understand how allometric plasticity evolves, it is necessary to study these selection pressures in detail – i.e., does the allometric scaling relationship of male sexual traits respond adaptively to different environmental cues for the expected future competitive environment? However, the evolution of allometric plasticity should be understood not just in terms of the selection pressures that might favour it, but also in terms of how allometric variation emerges from development. We may not even expect that allometries are food-independent, if we factor in the likely possibility that the energetic expenses of enlarged sexual trait production, relative to the total resource budget, depend on body size and growth rate.

Our observation of food-dependent allometric plasticity in *R. robini* adds to the limited number of recorded cases of allometric plasticity in male arthropod secondary sexual traits. Allometric plasticity is hypothesized to facilitate rapid local adaptation, adaptive divergence and speciation, if allometric plasticity occurs in the same direction as evolutionary change of trait allometries, and if plastic shifts in allometries coincide with release of cryptic genetic variation (Casasa & Moczek, 2019; Noble *et al*., 2019). To understand this potentially important evolutionary process, we should start looking for allometric plasticity in different species with conspicuous allometries in variable environments, and investigate whether the direction of allometric plasticity (Emlen, 1997; Knell *et al*., 1999; Moczek, 2002; Okada & Miyatake, 2010; Hardersen *et al*., 2011) generally coincides with evolutionary shifts in the allometry (Moczek *et al*., 2002; Moczek & Nijhout, 2003; Tomkins *et al*., 2011; Smallegange & Deere, 2014). Only then can we start to understand how development produces allometric variation, how or whether selection favours particular plastic responses of the scaling relationship, and eventually whether allometric plasticity contributes to evolutionary diversification.

## Supporting information

Analysis of measurement error

## Acknowledgements

We would like to thank Peter de Ruiter for providing feedback on earlier versions of this manuscript. IMS, KAS and FTR acknowledge a VIDI grant (No. 864.13.005) from the Netherlands Organisation for Scientific Research (NWO).

