## Supplementary material for "Nutrient-dependent allometric plasticity in a male-diphenic mite": Analysis of measurement error

**Supplementary Information: Analysis of measurement error**

**Methods**

We selected 7 adult male *R. robini* raised on 40 mg/mL (3 fighters, 2 scramblers) and on 2.56 mg/mL (2 scramblers) yeast solved in water. We photographed the mites ventrally and measured them to the nearest thousandth of a μm using a Zeiss Axiocam 105 color camera mounted on a Zeiss Stemi 200-C stereomicroscope, and ZEN 2 (Blue edition) software. We photographed each subject 10 times. Between photographs, we gently repositioned each subject on its back using a fine brush. We measured the maximum width of the idiosoma and the width of the left and right third femur. We measured the photographs not immediately after taking them, but after we were completely done with photographing our 7 subjects. We measured the photographs in blocks of 7, representing a single photograph of all 7 subjects, in pseudo-random order within blocks (the order in which subjects were measured was randomized within blocks) and among blocks (the order in which the photographs were collected into 7 blocks was randomized). Each block of measurements was recorded on a different spreadsheet, so we were blind to previous measurements.

To calculate the measurement error, we constructed a linear model for each of the three measurements, with the measurement as the dependent variable and individual ID as explanatory variable. We extracted the residuals and checked whether they were homogeneous across subjects using Bartlett's test of variance homogeneity. There was no evidence of variance heterogeneity across subjects in any of the measurements, so we proceeded by calculating the sample standard deviation of the residuals as our estimate of measurement error. If there would have been evidence of variance heterogeneity across subjects, we would have proceeded by investigating how the measurement error changed with body size.

**Results and discussion**

There was no evidence of variance heterogeneity in the model residuals for idiosoma width (Bartlett's test: $K^2 = 10.32$, df = 6, P = 0.112), right femur width (Bartlett's test: $K^2 = 8.57$, df = 6, P = 0.120) or left femur width (Bartlett's test: $K^2 = 4.84$, df = 6, P = 0.564), showing that the measurement error was the same for all subjects. The sample standard deviation of measurements (pooled for all subjects) was 3.3 μm for idiosoma width and 1.1 μm for both left and right femur width. Therefore, approximately 68% (normal distribution: $\mu \pm \sigma$) of the reported idiosoma width measurements are within 3.3 μm from the true value, and approximately 95% (normal distribution: $\mu \pm 2\sigma$) of the reported idiosoma width measurements are within 6.6 μm from the true idiosoma width value. Approximately 68% of the reported femur width measurements are within 1.1 μm from the true femur width value, and approximately 95% of the reported femur width measurements are within 2.2 μm from the true femur width value.
